# Assessment of asymmetric cell divisions in the early development of *Caenorhabditis elegans*

**DOI:** 10.1101/109215

**Authors:** Rolf Fickentscher, Matthias Weiss

**Affiliations:** Experimental Physics I, University of Bayreuth, D-95440 Bayreuth, Germany

**Keywords:** asymmetric cell division, embryogenesis, Caenorhabditis elegans, light sheet microscopy, SPIM, modeling

## Abstract

Asymmetric cell divisions are of fundamental importance for developmental processes, e.g. for the generation of founder cells. Prime examples are asymmetric cell divisions in the P lineage during early embryogenesis of the model organism *Caenorhabditis elegans*. However, due to a lack of quantitative data it has remained unclear how frequent unequal daughter cell sizes emerge in the nematode’s early embryogenesis, and whether these originate from sterical or biochemical cues. Using quantitative light-sheet microscopy, we have found that about 40% of all cell divisions in *C. elegans* until gastrulation generate daughter cells with significantly different volumes. Removing the embryo’s rigid eggshell revealed asymmetric divisions in somatic cells to be primarily induced by steric effects. Division asymmetries in the germline remained unaltered and were correctly reproduced by a model based on a cell-size independent, eccentric displacement of the metaphase plate. Our data suggest asymmetric cell divisions to be essential for establishing important cell-cell interactions that eventually fuel a successful embryogenesis.

**Summary statement:** About 40% of all cell divisions in early *C. elegans* embryogenesis are found to be asymmetric. A cell-size independent displacement of the mitotic spindle explains division asymmetries in the germline whereas the confining eggshell induces asymmetries of somatic cells.

## Introduction

Asymmetric cell divisions are of crucial importance for developmental processes, e.g. in the context of tissue or body axis formation (Betschinger and Knoblich, 2004, Rose and Gonczy, 2014). Many protein species that are involved in asymmetric cell divisions have been shown to be evolutionary conserved (reviewed, for example, in (Betschinger and Knoblich, 2004)), indicating that general mechanisms for asymmetry generation are utilized in different biological systems. Studies on the model organism *Caenorhabditis elegans* have been instrumental in this context due to its relative simplicity, its susceptibility to modern genetic and molecular-biological tools, and its optical transparency (see *www.wormbook.org* for an introduction). A plethora of (fluorescence) microscopy-based studies have, for example, revealed detailed insights into the first asymmetric cell division of the zygote (P_0_) and the concomitant creation of an *anterior-posterior* body axis (Boyd et al., 1996, Rose and Gonczy, 2014, Watts et al., 1996, Colombo et al., 2003, Grill et al., 2001, Goehring et al., 2011, Grill et al., 2003). Also a fair understanding of the associated formation of biochemical gradients, from Turing-like patterns (Goehring et al., 2011, Kondo and Miura, 2010) to condensation phenomena (Brangwynne et al., 2009), has been possible. Virtually all of these and similar studies have been focusing on the single-cell stage and the first, asymmetric cell division since monitoring dynamic intracellular events in the comparatively large P_0_ cell is straightforward.

In fact, although *C. elegans* has been studied as a model organism for several decades by now, cell division asymmetry has remained a rather vaguely defined term as it may describe purely biochemical or geometrical asymmetries, or the combination of both. Defining biochemical asymmetries of daughter cells necessarily requires the quantification of a nonuniform distribution of specific molecular markers and hence virtually all of such reported asymmetries are properly defined (see, for example, (Rose and Gonczy, 2014) for a comprehensive summary on biochemical asymmetries in the zygote). However, geometrical asymmetries, i.e. the emergence of two unequally sized daughter cells, have been studied in much less detail. Frequently utilized techniques like differential interference contrast (DIC) microscopy or even confocal microscopy have method-intrinsic limitations that hamper a thorough three-dimensional quantification, hence requiring simplifying extrapolations to arrive at approximate cell volumes (see (Galli and Morgan, 2016) for a recent example). Moreover, due to volume-conserving (blastomeric) division cycles, cell sizes in the early *C. elegans* embryo decrease rapidly, therefore amplifying the uncertainty about actual cell volumes. As a consequence, extrapolated cell volumes are quite error-prone and may not report reliably on geometrical asymmetries in cell division events.

Despite these limitations, it is well established that at least cells of the future germline, the so-called P lineage (cf. the embryo’s early lineage tree in Fig. 1A), undergo geometrically asymmetric divisions (Rose and Gonczy, 2014, Sulston et al., 1983). Yet, a thorough quantification of their (and other cells’) asymmetries has, to the best of our knowledge, not been done. As a consequence, it is neither clear how many geometrically asymmetric cell divisions beyond the P lineage occur until gastrulation nor is it known what causes them. Indeed, one may even ask why *C. elegans* has geometrically asymmetric cell divisions at all since a biochemical asymmetry might have been sufficient to run the proper molecular-biological developmental program.

**Figure 1.**
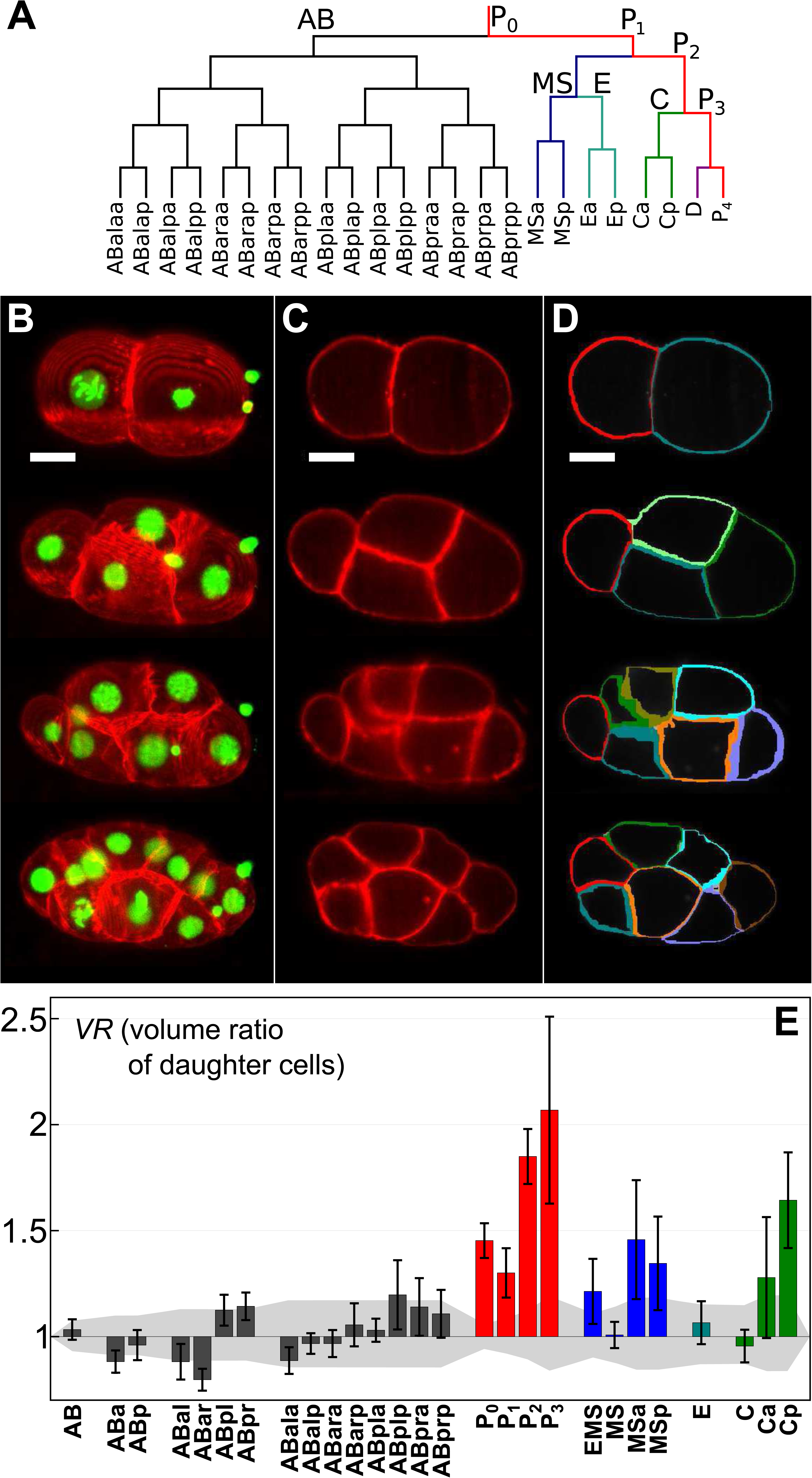
Division asymmetries in unperturbed *C. elegans* embryos. (A) Lineage tree of early *C. elegans* embryogenesis (prior to gastrulation). Different lineages are color-coded, the germline is highlighted in red. (B) Representative maximum-intensity projections of image stacks taken on early *C. elegans* embryos (strain OD95) with the plasma membrane and chromatin stained in red and green, respectively. Scale bar: 10 µm. (C) Single two-dimensional slices taken from the image stacks shown in A. (D) The corresponding membrane segmentation shows how well details of the plasma membrane are identified. (E) Volumetric ratio, *VR*, of daughter cells emerging from the named mother cell (median of *n=10* embryos with error bars indicating the standard deviation). Color-coding of lineages like in (A). The volume-dependent level of uncertainty for each cell (grey) quantifies the apparent division asymmetry that is attributed solely to segmentation errors (see Materials and Methods for a detailed definition). As a result, cells of the P, MS, and C lineages, but also few cells of the AB lineage show significant division asymmetries that are well beyond the level of uncertainty.

Here we have used selective plane illumination microscopy, SPIM, to address this topic (see, for example, (Hockendorf et al., 2012, Huisken and Stainier, 2009, Tomer et al., 2013) for introductory reviews on SPIM). Due to the gentle illumination via a light sheet, we were able to monitor the development of *C. elegans* embryos with and without an eggshell in three-dimensional detail up to gastrulation. A custom-made image segmentation approach enabled us to derive volumes and division asymmetries from these raw data. As a result, we observed that about 40% of all cell divisions before gastrulation are significantly asymmetric with many of these events being enhanced by sterical forces from the confining eggshell. For predominantly biochemically governed asymmetric cell divisions, i.e. for the P lineage, we were able to predict the degree of asymmetry via a simple model that relies on a cell-size independent, eccentric displacement of the mitotic spindle.

## Results and Discussion

### About 40% of all cell divisions until gastrulation are asymmetric

According to the literature, asymmetric cell divisions have been observed for cells of the P lineage (P_0_,…, P_3_) and EMS (Rose and Gonczy, 2014, Schierenberg, 2006, Sulston et al., 1983). These cell divisions coincide with the emergence of so-called founder cells (AB, MS, E, C, D, P_4_) that establish new lineages (Fig. 1A). All other cell divisions until gastrulation are typically interpreted as being symmetric with respect to daughter cell sizes. In fact, data on volumetric asymmetries of daughter cells are mostly qualitative or extrapolated from two-dimensional imaging techniques rather than reporting faithful three-dimensional quantifications.

In order to obtain more quantitative insights into the amount and degree of volumetric asymmetries in cell divisions during early embryogenesis of *C. elegans*, we used a custom-made SPIM setup (Fickentscher et al., 2013, Fickentscher et al., 2016, Struntz and Weiss, 2016) and a custom-written segmentation approach (see Materials and Methods for details). The chosen worm strain (OD95) stably expressed fluorescent markers for histones (H2B∷mCherry) and the plasma membrane (PH(PLC1δ1)∷GFP), hence facilitating three-dimensional imaging and volume rendering during early embryogenesis. Representative examples of images and segmentation results are shown in Fig. 1B-D.

Using this approach we were able to identify all cell division events and to quantify all cell volumes until the onset of gastrulation via temporally resolved three-dimensional image stacks. Using these data as experimental input, we calculated for each mother cell the volume ratio of daughter cells, *VR=V_1_/V_2_*. For somatic cells we used the more posterior cell for *V*_*2*_, for germline cells always the (smaller) new germline cell volume was used for *V*_*2*_. To explore whether a division was significantly asymmetric, i.e. whether the value of *VR* deviated sufficiently from unity, we defined a level of uncertainty that arises solely from segmentation errors (see Materials and Methods for details). Roughly speaking, relative deviations of daughter cell volumes by 10% or more indicated a significant volumetric asymmetry in the respective cell division. The resulting data and their significance rating are shown in Fig. 1E.

As expected, all cells of the P lineage showed a significant division asymmetry, albeit with markedly different *VR* values (Fig. 1E). Also EMS was found to divide asymmetrically, although the asymmetry was less than for any cell of the P lineage. Surprisingly, also other cells at this early stage, namely MSa, MSp, Ca, and Cp, showed significant asymmetries with daughter cells differing by 30-60% in volume. In contrast, E, MS, and C divided almost perfectly symmetrical like most cells from the AB lineage. Yet, even some cells from the AB lineage showed a significant but borderline asymmetry, especially ABar.

Thus, our data indicate that altogether about 40% of all cell divisions until gastrulation in *C. elegans* embryos feature a significant volumetric asymmetry.

### Geometrical constraints induce asymmetric divisions

Since, to our knowledge, a biochemical asymmetry in the cell division of MSa, MSp, Ca, and Cp has not been described, we hypothesized that at least some of the observed volumetric asymmetries might arise from geometric constraints during the respective cell divisions rather than being governed by biochemical cues. Indeed, geometrical constraints and physical forces have been seen to have a significant impact on the positioning of cells in the early embryo (Fickentscher et al., 2013, Fickentscher et al., 2016) but also in the context of tissue organization and wound healing (Puliafito et al., 2012, Streichan et al., 2014), making this hypothesis an attractive option.

In order to test our hypothesis, we sought to decrease possible mechanical constraints by removing the chitin-based eggshell, while leaving the inner, more flexible vitellin layer intact to maintain the embryo’s integrity (see Materials and Methods). As expected, in the absence of the eggshell the intact vitellin layer lead to a rather compact arrangement of the blastomeres that was often, but not always, similar to wild-type embryos (Fig. 2A-C). The associated volume ratios (Fig. 2D) and the direct comparison to untreated embryos (Fig. 2E) highlighted that the P lineage (P_0_ to P_3_) as well as Ca did not change markedly. This suggests that their volumetrically asymmetric divisions are a consequence of an internal biochemical asymmetry. Almost all other volume ratios showed a clear tendency to decrease, indicating that sterical forces induced by the confining eggshell cause at least partially these asymmetries. In particular, EMS and the MS lineage showed markedly reduced asymmetries, making them almost as borderline as the somatic outlier ABar. Cp showed a significantly reduced value of *VR* while still dividing in significantly asymmetric fashion, whereas Ca was almost unchanged.

**Figure 2.**
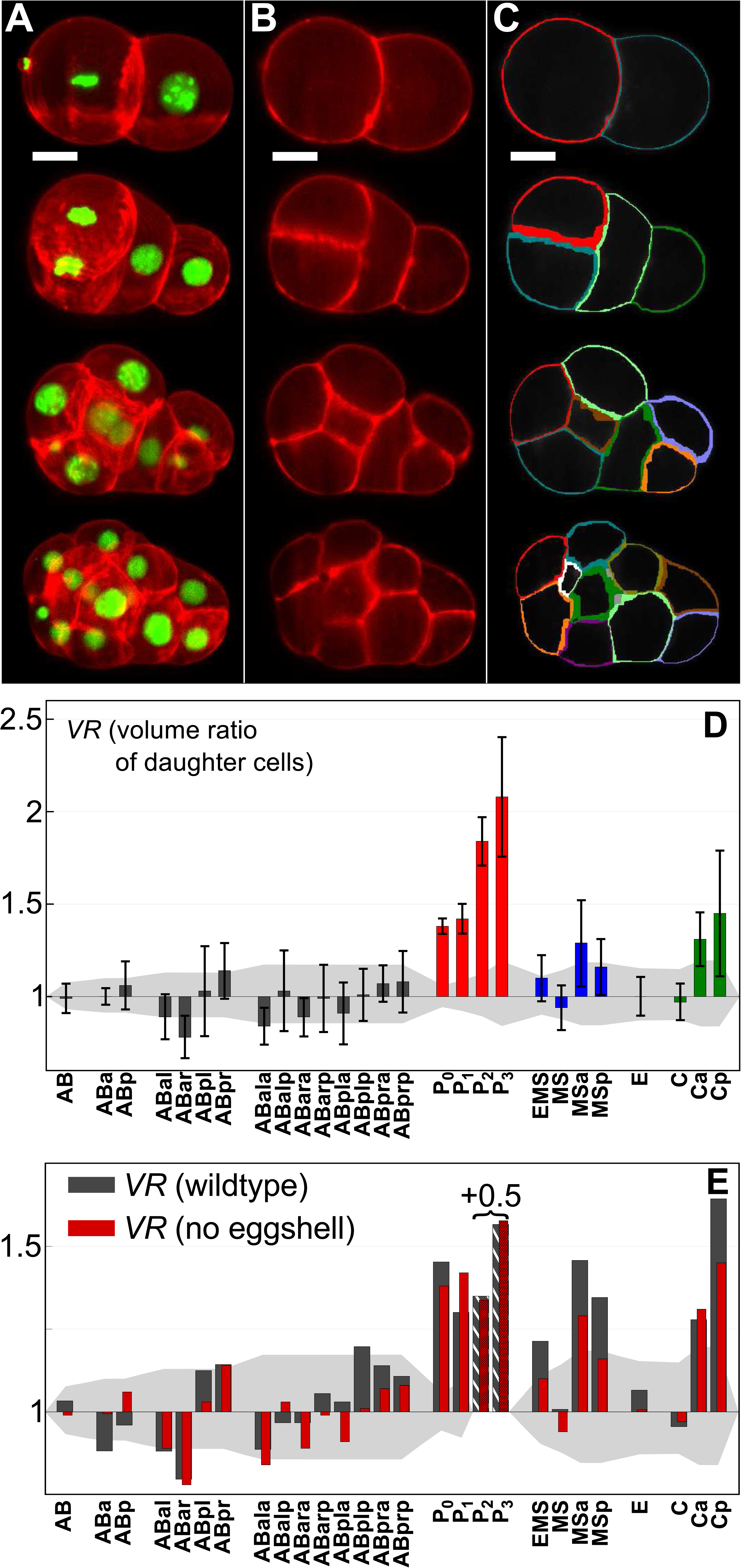
Division asymmetries in *C. elegans* embryos lacking the eggshell. (A) Representative maximum-intensity projections of early *C. elegans* embryos after removing the eggshell but leaving the vitellin layer intact (red, green: plasma membrane, chromatin). Scale bar: 10 µm. (B,C) Single two-dimensional slices taken from the image stacks shown in A, and corresponding membrane segmentation. (D) Volumetric ratio, *VR*, of daughter cells emerging from the named mother cell (median of *n=11* embryos, error bars indicate the standard deviation) with the volume-dependent level of uncertainty. Color-coding as in Fig. 1E. (E) Comparison of the median values without (red) and with (grey) an intact eggshell. Please note that bars for P_2_ and P_3_ have been reduced by 0.5 for better visibility. While asymmetries in the germline are preserved, most somatic cells tend to decrease their level of asymmetry upon removal of the eggshell.

Thus, asymmetric divisions in the P and C lineages are well preserved even with softened geometric constraints while the asymmetry in EMS and in the MS lineage seem to rely predominantly on sterical forces imposed at least indirectly by the eggshell.

### A cell-size independent displacement of the mitotic spindle quantitatively explains division asymmetries in the germline

Inspired by previous work on the first cell division in *C. elegans* embryos, in which a pronounced shift of the mitotic spindle apparatus along the AP-axis has been identified as major cause for an asymmetric cell division of P_0_ (Colombo et al., 2003, Grill et al., 2001), we hypothesized that also subsequent asymmetric cell divisions in the germline are driven by a displacement of the spindle’s center of mass. In particular, we wondered to which extent a shift of the mitotic spindle could quantitatively explain the experimentally observed volumetric ratios *VR* of daughter cells in the post-zygote germline. For this analysis, we deliberately excluded P_0_, since several molecular players that influence an eccentric spindle displacement in P_0_ from the anterior side (Galli et al., 2011, Panbianco et al., 2008) are segregated into the AB cell during the first cell division. Hence, they are unlikely to play a major role in subsequent cell divisions in the germline, rendering the first division a special case (see also discussion below). Moreover, we have used data from embryos without eggshell for the subsequent analysis since these display asymmetric divisions without the influence of eggshell-induced cues; we did not observe significant differences when using data from untreated embryos.

For simplicity we assumed the mother cell to be spherical (radius R_0_), and we asked how the volumetric ratio would be if the daughter cells emerged from a spindle that was displaced by an increment Δx away from the cell center (Fig. 3A, inset). Given that the position of the metaphase plate defines the plane of cytokinesis (Rappaport, 1971), the spherical mother cell would be split approximately into two spherical caps with heights R_0_+Δx and R_0_-Δx, yielding volumes V_1_=π/3·(R_0_+Δx)^2^·(2R_0_-Δx) and V_2_ = π/3·(R_0_-Δx)^2^·(2R_0_+Δx), respectively. The mathematically simplest assumption for this division scheme would be a constant shift Δx that does not depend on the size of the mother cell.

**Figure 3.**
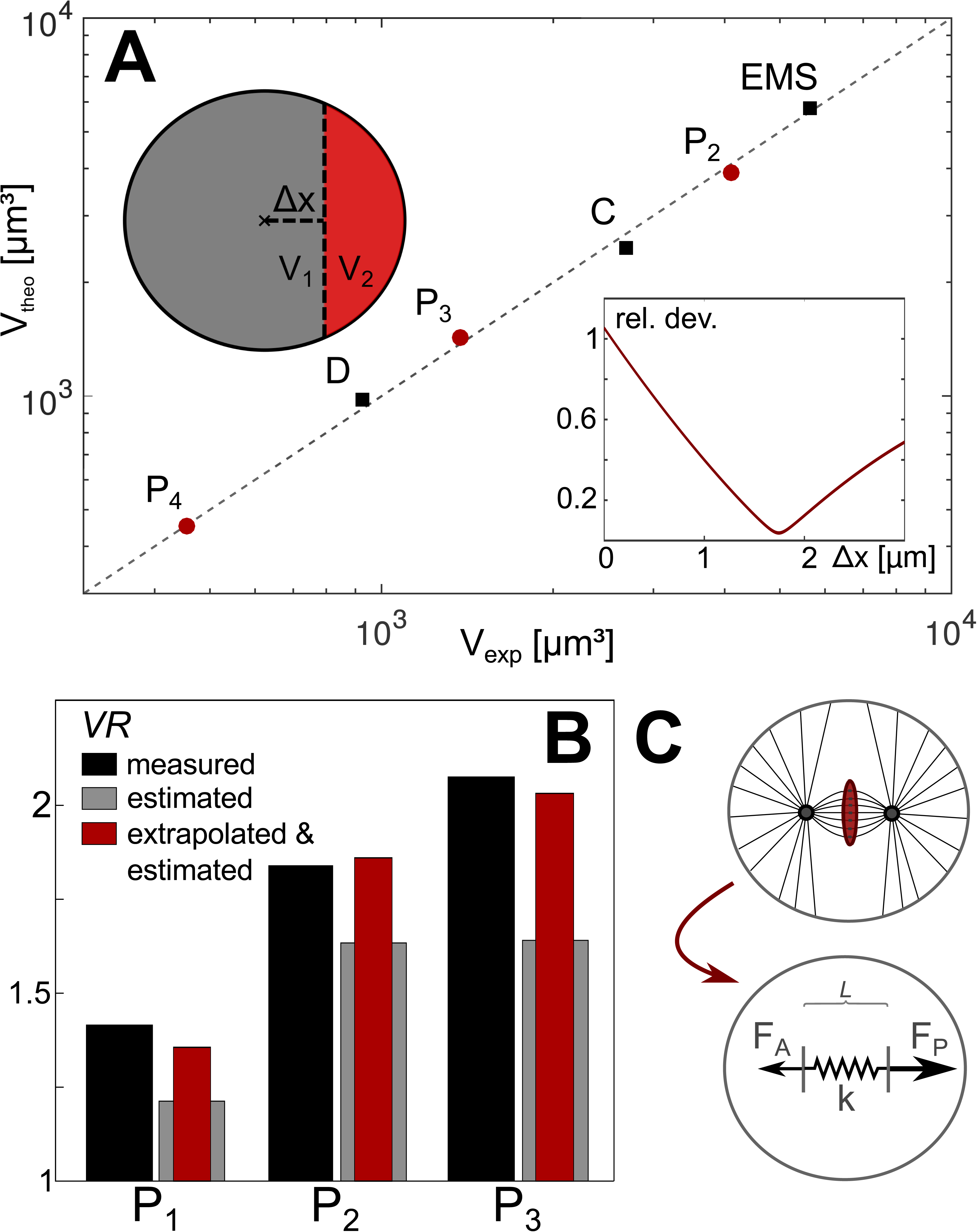
Comparison of experimental findings and model predictions. (A) Modeling cells as spherical entities and allowing for an off-center division into spherical caps (cf. inset upper left) predicts well the experimentally observed cell volumes V_exp_ (main plot). Using the volume of P_1_ as sole input, volumes of EMS and P_2_, and from this volumes of P_3_ and C, and finally P_4_ and D were deduced assuming a constant shift Δx of the division plane. Predicted values V_theo_ matched best the experimentally found ones, when choosing Δx=1.75µm (see inset lower right for the relative deviations when varying Δx). (B) Division asymmetries *VR* predicted for P_1_, P_2_, and P_3_ on the basis of the last image stack that shows an unambiguous metaphase (grey bars) follow the experimental results for the daughter cells (black bars) but consistently underestimate the asymmetry. In fact, using these image stacks and dissecting segmented cells into two caps via a division plane through the metaphase plate reveals a median spindle shift of only 1.35µm (see main text for details). Accounting for an additional, unmonitored spindle shift by approximately 450nm during the lag period between consecutive image stacks (see main) and repeating the dissection scheme the extrapolated asymmetries (red bars) show a favorable agreement with our experimental data. (C) Spindle displacement by a constant offset Δx can be rationalized by assuming constant forces F_A_ < F_P_ that pull the spindle towards the anterior and posterior end of the cell, respectively. As a result, the spindle is stretched and its center of mass moves into the posterior direction. Stress resistance of the spindle is modeled via a passive Hookean element (spring constant k, resting length L_0_) until a maximum extension is reached and the spindle ruptures at the onset of anaphase. See main text for details.

To compare this naïve approach with our experimental data, we used the measured volume of P_1_ from which we extracted the apparent cell radius via R_0_=(3V/4π)^1/3^. Then, we iteratively predicted from this single experimental input the volumes *V*_*theo*_ of all subsequent daughter cells along the lineage tree (P_2_, EMS, P_3_, C, P_4_ and D): Volumes of EMS and P_2_, i.e. V_1_ and V_2_, were derived via the spherical-cap scheme outlined above, using R_0_ of P_1_ as input. Assuming P_2_ to be spherical again, we extracted its apparent radius from the predicted volume and repeated the division scheme until all volumes had been determined. The predictions we got from this procedure showed a remarkably good agreement with the experimentally observed volumes of daughter cells, V_exp_, when setting Δx≈1.75μm (Fig. 3A). Using instead a shift Δx that depended on the mother cell size did not capture the experimental data.

Thus, a cell-size independent shift of the mitotic spindle by approximately 1.75μm can quantitatively explain all experimentally observed volumetric division asymmetries in the post-zygote germline.

Next we sought to obtain experimental support for this simple approach and its prediction of a cell-independent displacement of the mitotic spindle by Δx≈1.75μm during asymmetric division events in cells P_1_-P_3_. In contrast to the distinct first cell division, division axes of cells P_1_-P_3_ do not necessarily lie in a single imaging plane, which virtually eliminates the possibility to determine the spindles’ shift via very rapid two-dimensional imaging. We therefore utilized our three-dimensional image stacks, acquired with a moderate time resolution, as a proxy.

In particular, we exploited the last image stacks in which P_1_, P_2_, and P_3_ showed an unambiguous metaphase pattern. Combining segmented image stacks and tracking data of chromatin, we determined the respective cell’s center of mass and the position and orientation of its metaphase plate (see Materials and Methods for details). Assuming the plane of division to coincide with the metaphase plate, we determined the distance of the cell center from the metaphase plate along its surface normal as an estimate for the eccentric displacement of the spindle. The median of all values collected for P_1_-P_3_ in 11 embryos was 1.35μm which is clearly nonzero but somewhat lower than the predicted value, Δx≈1.75μm. We attributed this difference to the fairly long lag time of 30s between successive image stacks, i.e. even after the last stack with a metaphase phenotype the spindle could still be on the move for up to 30s. Based on data acquired for P_0_ (Grill et al., 2001) a peak velocity in the range of 40nm/s can be expected for the spindle motion. Assuming that all spindles in the post-zygote germline move with a somewhat lower, average velocity of 30nm/s (see below for a justification) and estimating that the onset of cytokinesis happens 15s after the image stack has been taken, an unmonitored distance of 15s×30nm/s=450nm should be taken into account to extrapolate the typical shift of the spindle in cells P_1_-P_3_. The result, a shift by Δx≈1.35μm+450nm=1.80μm, is in favorable agreement with our prediction derived via the division scenario into spherical caps.

This extrapolation is further corroborated by the volumetric asymmetries, *VR*, determined from the very same segmented image stacks: Using the metaphase plate to determine the future division plane, all voxels of cells P_1_-P_3_ were sorted into putative daughter cells (see Materials and Methods), i.e. real cells were dissected through the metaphase plate into slightly deformed spherical caps. Values of *VR* determined via this procedure indeed followed the trend of the experimental data for fully developed daughter cells (Fig. 3B), yet consistently underestimated the asymmetry. Shifting the putative division plane by 450nm along its surface normal to account for the spindle movement between successive image stacks, we obtained a favorable agreement between the estimated and experimentally determined asymmetries (Fig. 3B).

Thus, the experimentally determined shift of the metaphase plate in cells P_1_-P_3_ confirms the reasoning of a cell-size independent eccentric position of the spindle before asymmetric division events.

### A simple model explains the constant spindle displacement

The above results raise the question of how cells actually ensure a size-independent spindle displacement to an eccentric position before entering anaphase. To arrive at a meaningful model, we started from experimental observations in the zygote: In P_0_, the anterior spindle pole is tethered and remains in a fixed position until the spindle is fully assembled (Labbe et al., 2004). Upon release of the tethering, the metaphase plate starts to displace to the posterior with a constant velocity (Labbe et al., 2004). While the anterior spindle pole is displaced only slightly to the posterior (Labbe et al., 2004), the posterior spindle pole is displaced significantly stronger (Colombo et al., 2003). Hence, concomitant to its displacement the spindle is stretched along its migration path. Moreover, laser ablation experiments in P_0_ have revealed that astral microtubules provide the pulling forces for the metaphase plate’s displacement (Colombo et al., 2003, Grill et al., 2001, Grill et al., 2003, Labbe et al., 2004). These pulling forces are generated at the cell cortex, at which astral microtubules are in contact with force generator complexes that involve Gα, GPR-1/2, LIN-5 and dynein (see (Rose and Gonczy, 2014) for a recent comprehensive review). A net force ratio of ~1.5 towards the posterior is achieved in P_0_ by locally increasing pulling forces on the posterior cortex (Colombo et al., 2003, Grill et al., 2001, Grill et al., 2003) but also by decreasing forces on the anterior cortex (Galli et al., 2011, Panbianco et al., 2008). Using RNA interference to switch both poles to the same force generator phenotype lead to a vanishing net force on the spindle (Grill et al., 2001), and consequently these embryos lacked the native spindle displacement towards the posterior end. When both poles were forced to assume an ‘anterior’ force generator phenotype, mitosis even was stalled in late metaphase due to a too low absolute force that was insufficient to initiate a rupturing of the spindle.

Combining these observations with our results, we can formulate a simple one-dimensional model for achieving a cell-size independent displacement of the spindle that precedes an asymmetric division (Fig. 3C): Forces acting on the anterior and posterior spindle poles, F_A_ and F_P_, do not depend on the distance to the cell cortex as microtubules only transmit pulling forces that are created at the cortex. Due to a low residence time of each microtubule at the cortex, about 1-2s (Kozlowski et al., 2007), microtubules also do not contribute a memory-driven restoring force. Therefore, F_A_ and F_P_ can be modeled as constant forces. The spindle does not provide active forces for its displacement but needs to oppose the net stress F_p_+F_A_ applied via the spindle poles. For simplicity, we model it as a Hookean element with spring constant *k* and equilibrium length *L*_*0*_. Since initial spindle lengths at early metaphase are almost cell-size independent at these early stages of embryogenesis (Hara and Kimura, 2009), we can assume *L*_*0*_ to be approximately the same for all P cells. Upon stretching this spring beyond a limit *L*_*0*_*+s*_*max*_, the spindle is assumed to rupture. This assumption is based on the observation that pulling on both spindle poles with only the anterior force magnitude is insufficient to go from metaphase to anaphase, whereas bidirectional pulling with the posterior force magnitude allows for spindle rupturing and (symmetric) cell division (Grill et al., 2001). Due to the low Reynolds number in cell biology problems (Phillips, 2013), the motion of the spindle and/or its poles can be described in the overdamped limit, i.e. we can neglect all inertia terms.

For convenience, we express the equations of motion in terms of the distances x_A_ and x_P_ that the anterior and posterior spindle poles assume over time with respect to their initial position. Initially, the two poles are separated by the equilibrium length *L*_*0*_ of the unstressed spindle, i.e. x_A_(t=0)=x_P_(t=0)=0. Upon releasing the tethering at time t=0, the forces F_A_<F_P_ act on the spindle poles and the spindle is stretched by a distance s=x_P_-x_A_. Hence, the equations of motion read for t>0:

-*γ*·dx_A_/dt + *k*(x_P_ – x_A_) – F_A_ = 0 and -*γ*·dx_P_/dt - *k*(x_P_ – x_A_) + F_P_ = 0,

with *γ* denoting the effective friction coefficient for the spindle poles. Solving these coupled differential equations, one obtains x_MP_(*t*) = [x_A_(t)+x_P_(t)]/2 = (F_P_-F_A_)·*t*/(2*γ*) for the position of the metaphase plate, and s(*t*)=(F_P_+F_A_)/(2*k*)·(1-exp(-2*kt*/*γ*)) for the extension of the stressed spindle. Upon reaching a maximum extension *s*_*max*_, the spindle ruptures and cytokinesis is initiated. The associated instant of time, *T*, is determined via the equation *s*_*max*_=s(T)=(F_P_+F_A_)/(2*k*)·(1-exp(-2*kT*/*γ*)) from which one can infer the maximum travel distance of the spindle, Δ*x*=x_MP_(*T*)= (F_P_-F_A_)·*T*/(2*γ*). Since neither the forces F_A_ and F_P_ nor the spindle parameters dependent on cell size, this model predicts a constant displacement of the mitotic spindle.

It is worth noting that the spindle displacement explicitly depends on the ratio of anterior and posterior pulling forces, F_A_/F_P_. In fact, adapting the aforementioned approach of spherical caps to the ellipsoidal zygote, a displacement of ~3μm was needed to explain the asymmetry ratio *VR* for P_0_, which is significantly larger than the value Δx≈1.75μm for P_1_-P_3_. This apparent discrepancy can be rationalized when taking into account that in P_0_ several molecular agents are localized in the anterior domain of the cortex where they contribute to a lowering of pulling forces towards the anterior (Galli et al., 2011, Panbianco et al., 2008). After the first cell division, these very proteins are segregated into the somatic cell AB and hence are lost for the future germline. As a consequence, these players will not be available to reduce F_A_ in cells P_1_-P_3_, hence decreasing the ratio F_A_/F_P_ in these cells. This leads to a slower spindle displacement in P_1_-P_3_ in comparison to P_0_ (as assumed when estimating the unmonitored spindle shift by 450nm), whereas the spindle-internal stress builds up more rapidly. Therefore, upon reaching its maximum extension s_max_, the spindle has travelled a smaller distance. Following this reasoning, cells P_1_-P_3_ are predicted to display a smaller displacement Δx than P_0_, in favorable agreement with experimental observations.

Finally, one may wonder why nature has chosen to equip *C. elegans* with volumetrically asymmetric cell divisions during early embryogenesis, as biochemical asymmetries could have been fully sufficient. While cell sizes seem to have little influence on the positioning of cells until gastrulation (Fickentscher et al., 2013, Fickentscher et al., 2016), the number of cell-cell contact areas certainly depends quite strongly on the surface area of cells. We therefore speculate that distinct division asymmetries cause, or at least support, the formation and/or prevention of cell contact areas to achieve a wiring diagram of cells that can fuel a successful embryogenesis.

## Author contributions

R.F. performed and analyzed all experiments and wrote all related software; M.W. conceptualized the study and designed experiments; R.F. and M.W. wrote the manuscript.

## Materials and Methods

### Sample preparation and imaging

For imaging we used *C. elegans* strain OD95 in which the plasma membrane and histones are fluorescently labeled (PH(PLC1δ1)∷GFP, H2B∷mCherry). Worm culture and preparation of untreated embryos for SPIM imaging was done as described before (Fickentscher et al., 2013, Fickentscher et al., 2016, Struntz and Weiss, 2016). Removal of the eggshell was done similar to previous approaches (Edgar and McGhee, 1988): Zygotes with visible pronuclei were chosen from dissected gravid worms. These eggs were placed on a coverslip and immersed in approximately 30μl of NaOCL solution (3% Na) for two to three minutes. Subsequently they were washed three times with M9 buffer (Shaham, 2006) to remove all of the NaOCL solution before pipetting 25μl Chitinase solution onto them. The solution was prepared by dissolving 5 units of Chitinase from Streptomyces (Sigma) in 2ml of sterile egg buffer (Shaham, 2006).

Removal of the eggshell took roughly 10 to 15 minutes. The remaining vitellin layer was left intact. When the eggshell was not visible any more, embryos were transferred rapidly to the SPIM setup for immediate imaging (starting in most cases during mitosis of the zygote). Embryos adhered to the plain, untreated glass surface without the need for Poly-L-lysine or other mounting agents. During imaging, unperturbed embryos were immersed in water, embryos without eggshell in M9 buffer. Imaging was performed with a custom-made dual color SPIM setup as described before (Fickentscher et al., 2013, Fickentscher et al., 2016, Struntz and Weiss, 2016). For long-term imaging of wildtype and eggshell-free embryos, full dual-color stacks, consisting of 50 individual layers with a spacing of √2 μm, were taken every 30s for a total time of three hours (i.e. 360 stacks).

### Segmentation, image analysis, and evaluation

Tracking of nuclei via H2B∷mCherry was done as described before (Fickentscher et al., 2013). We generally tracked at least until the embryo consisted of 44 cells and included a manual correction step to account for potential errors.

Three-dimensional segmentation of cell membranes from PH(PLC1δ1)∷GFP images required a refined approach to account for SPIM-inherent shadowing effects. These arise from absorption and scattering events at bright structures when being illuminated by the light sheet, i.e. some shadowing is observed behind such structures. As a result, image segmentation of membrane-labeled embryos via global filtering and thresholding operators was not reliable. We have therefore developed a novel segmentation algorithm that is based on growing a seed region in each cell: The goal of the segmentation is a division of a three-dimensional image of an embryo with *n* cells into *n+1* regions, with each voxel of the image stack being uniquely assigned to one of the cells or to none (= outside of the embryo). After an initial box-filtering (kernel size 5×5×1 voxels) the background intensity of the image, i.e. anything outside of the embryo, is set to zero via global thresholding. Then, a seed is placed inside each cell, either manually or by using voxels that have been identified during the tracking of nuclei. During the segmentation process, these *n* seeds are grown simultaneously and iteratively. Boundaries of each seed are computed by eroding the region of voxels belonging to the seed with the smallest possible kernel (3×3×3 voxels) and subtracting this result from the original region. Boundary voxels therefore share one face, edge, or corner with voxels outside of this seed’s region. Next, for each of these boundary voxels one neighbor outside the region is chosen randomly and both voxel intensities are compared. If the outside voxel’s intensity is larger than the boundary voxel’s intensity (*F*_*out*_*≥qF*_*boundary*_), the outside voxel is added to the region unless it belongs already to another seed’s region. The multiplier *q≈0.97…0.99* is introduced to compensate for noise, and it needs to be chosen carefully for each image or image series. This procedure is carried out for all cells/ seed regions prior to the next iteration. Aiming at short processing durations, a total of *N=300/log(n+1)* iterations were performed initially on a downscaled version of the image stack. After upscaling to the original size, 40 additional iterations were performed.

This scheme leads to a local expansion of each seed until it collides with another region or when its boundary arrives at a significant drop in voxel intensity. The latter typically occurs at the cytoplasm-membrane interface, i.e. the boundary of each region becomes a faithful representation of the plasma membrane. Minor artifacts (stray pixels etc.) are removed after the iteration process by opening and closing operations with a kernel of 15×15×3 voxels. Results were finally controlled manually stack by stack to ensure a high segmentation quality. Results obtained with this algorithm provided us with data of cellular volumes (and shapes) of unprecedented precision.

From the segmentation process, the number of cells, *n*, and, the number of voxels of each cell, *m*_*i*_ (*i=1,…, n*), is known for each embryo at each instant of time. From this, each cell’s volume was determined as the product *V*_*i*_*=L*_*x*_*L*_*y*_*L*_*z*_*m*_*i*_ with *L*_*x*_*=L*_*y*_*=*0.16µm being determined via the objective and the camera sensor, and *L*_*z*_=√2μm the spacing between two consecutive optical sections within an image stack. Volumes do not show any significant changes during the cell cycle, i.e. a cell’s volume can be assumed constant (data not shown). For our analysis, we used the median value of the obtained time series of cell volumes to suppress few possible outliers in the scheme. As a result, the sum of volumes of the daughter cells generally deviated from the mother cell’s volume by less than 3%, indicating reliable segmentation results throughout the whole image series.

Volume ratios of somatic daughter cells were defined as the volume of the more anterior cell divided by the volume of the more posterior cell, e.g. *VR*_*AB*_*=V*_*ABa*_*/V*_*ABp*_. For germline cells always the smaller volume of the new germline cell was used in the denominator, e.g. *VR*_*P3*_*=V*_*D*_*/V*_*P4*_. The level of uncertainty due to segmentation errors, i.e. the minimal ratio *VR* that reports a significant division asymmetry, was determined as follows: The main source of error during volume determination originates from the decision whether or not an additional layer of voxels around the already detected volume is considered while segmenting a cell, i.e. if the region is expanded even further or not. Assuming all cells to be spherical with radii being determined by the cell volume, *R* = (3*V* / 4π)^1/3^, we can express this additional volume ΔV in analytical terms. We first note that due to L_z_ >> L_x_ we have to separately consider the bottom and top slice in the (sub)stack containing the respective cell, i.e. we have two contributions, ΔV_1_ and ΔV_2_, from the inner layers and the two spherical caps in the top and bottom layer, respectively. We reasoned that the true cell boundary will, on average, bisect the thickness of the top and bottom layers (cf. sketch in Fig. S1), i.e. top and bottom caps have a height h=L_z_/2 and a squared in-plane radius r^2^=R^2^-(R-h)^2^ yielding a volume contribution 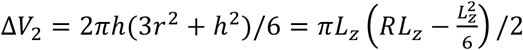. The contribution from the inner layers, where additional voxels have an average width L_x_, is simply the volume of a spherical shell with thickness L_x_, reduced by the volume of the two caps that are situated in the top and bottom layer, ∆*V*_1_ = 4*πR^2^L_x_* − 2*πRL_z_L_x_*. Since any of these additional volumes will contribute only by chance, adding or not adding these voxels has equal probability 50%, the average volume that is added or not considered amounts to ΔV=(ΔV_1_+ΔV_2_)/2. Hence, even a symmetric division into daughter cells with volume *V*_*sym*_ can lead to an apparent asymmetry *VR*_*max*_=(*V*_*sym*_+ΔV)/(*V*_*sym*_-ΔV)>1 or *VR*_*min*_=(*V*_*sym*_*-*∆V)/(*V*_*sym*_+∆V)<1, depending on which daughter cell determines the ratio’s (de)nominator. Based on this reasoning, we only deemed measured asymmetries as significant when they exceeded this mother-cell volume-dependent uncertainty range.

To estimate the spindle displacement in late metaphase and extrapolate the future division plane, we used three consecutive stacks (named S_1_,…, S_3_) after the last image stack that showed an unambiguous metaphase phenotype (named S_0_): From stack S_0_, the center-of-mass position of the metaphase plate, **r**_0_, and of the entire cell, **c**, are known from tracking the histone stain (H2B∷mCherry) and segmenting the plasma membrane stain, respectively. Positions **r**_1_ and **r**_2_ of daughter-cell chromatin assemblies are known for S_1_,…, S_3_ from tracking the histone stain (H2B∷mCherry). Normalizing and averaging the vector **r**_2_ – **r**_1_ over stacks S_1_,…, S_3_ yielded a robust estimate for the surface normal of the future division plane, **d**. The distance by which the metaphase plate had been shifted to an eccentric position in stack S_0_ was then determined as the scalar product Δx = **d·(c-r**_0_). Then, the position of the metaphase plate was split into two artificial points **q**_1_ and **q**_2_ along the direction of **d** with a small separation of 10nm (<<L_x_), i.e. **q=r**_0_± **d**·5nm. Based on the shortest distance to these points, all voxels of the cell were then classified to belong to **q**_1_ or **q**_2_, and the resulting volumes V_1_ and V_2_ were used for Fig.3A. Indeed, this scheme splits the volume of the mother cell into two parts along a plane (perpendicular to **d**) that runs through the last metaphase plate position, **r**_0_. When extrapolating the unmonitored spindle movement between stacks S_0_ and S_1_, the same approach was used with points **q**_1_ and **q**_2_ being shifted by **d**·450nm.

All evaluation codes written in Matlab are available upon request.

## Acknowledgment

Financial support from DFG grant WE4335/3-1 is gratefully acknowledged. Worm strains were provided by the Caenorhabditis Genetics Center, which is funded by the NIH Office of Research Infrastructure Programs (P40 OD010440). The authors declare no competing financial interests.

